# Proximo-distal muscle modulation as a function of hand orientation in a reach-and-grasp task

**DOI:** 10.64898/2026.03.27.714710

**Authors:** Chambellant Florian, Hilt Pauline, Cronin Neil, Thomas Elizabeth

## Abstract

The aim of this study was to improve our understanding of muscle contractions in the arm as a function of hand orientation for grasp. While there have been several reports on arm kinematics for reach and grasp movements, little has been done at the muscular level. To this end, we analyzed the modulation of shoulder, elbow and hand muscles for a reach and grasp task involving a target in either horizontal or vertical orientation. We hypothesized that unlike what has been observed for kinematics, at the muscular level we would see less correlation between the three muscle groups. A decoding approach with Machine Learning revealed adaptation patterns that were not visible using classical methods. Reach-and-grasp has traditionally been treated as being made of two components – the reach and the grasp components. Our dynamic decoding approach revealed a more complex picture with very different dynamics in the shoulder and elbow muscle groups during reach. All muscle groups showed peak capacity for predicting hand orientation before the start of grasp and followed the ubiquitous proximo-distal organization. The patterns of muscular modulation for hand orientation were strongly perturbed by the ‘eyes closed’ and ‘slow movement’ conditions, potentially decreasing the available degrees of freedom for adaptation.

## Introduction

The aim of this project was to improve our understanding of the reach and grasp task with the target at different orientations. While there have been several studies on the kinematics of the movement [1], [2], [3], [4], [5], [6], [7], very few studies have analysed the muscular adaptation for this target change. In particular, we wanted to investigate the differences in the activations of the shoulder and elbow muscle groups.

A majority of the discussions around reach-and-grasp has focused on the question of whether the reach component and the grasp component of this task are to be considered as a unified whole with a common controller or whether they operate with independent controllers [8], [9]. On the side of independent control, several researchers demonstrated that extrinsic properties of the target such as target distance, change the kinematic characteristics of reach without changing the nature of the grip component [8], [10]. Jeannerod (1981) demonstrated that key characteristics of grip such as time to peak grip aperture occurred at approximately 75% of movement duration irrespective of target distance. Furthermore, the two visuomotor channels of the nervous system, the dorsal and ventral streams [11], [12], [13] were proposed as a possible substrate for separating these two components [14], [15]. It was suggested that the dorsal stream of the visuomotor pathway carries information pertaining to the reach or transport component, hence being more influenced by the extrinsic aspects of the task such as distance and position of the target. Grip formation on the other hand was thought to be more controlled by the ventral stream and determined by the intrinsic properties of the target such as size or shape.

While an independent configuration of these two components has been observed using kinematics, many others have shown the two to have an influence on each other, i.e. perturbation of one component alters the other. For example, an unexpected change in the location of a target was also found to alter grip characteristics [16], [17]. Alterations in the requirements of grip were found to modify the kinematics of reach. An example of this would be the influence of object size on reach [18]. More specifically, with regards to the topic of the current investigation which centres around the influence of object orientation, Desmurget et al [3], [5] showed that wrist trajectories during the reach phase of reach-and-grasp are altered by object orientation.

While the debate has centred around the interaction of these two components – reach and grasp, there are in fact two components involved in reach itself – movement around the shoulder as well as elbow joint. Other than the well known correlations between the kinematic angles of the arm during simple reach [19], [20], various kinematic studies have also shown a good degree of correlation between these two components during reach for reach-and-grasp [3], [5], [21]. The elbow joint however, is closer to the hand effector for grip than the shoulder. It would be natural to expect some differences in the contributions of these two joints for hand orientation. Here, we put forward the hypothesis that these differences will be much more visible at the muscular level. Indeed, the muscular system being highly redundant, it is possible for muscles to have different patterns of underlying activity even as the kinematics of the movement are identical [22], [23].

In the current investigation, we also studied the effects of alterations in sensory feedback for muscular adaptation to object orientation. Questions on the role of feedforward and feedback control are very active in movement studies [24], [25], [26], [27], [28]. The domain of reach and grasp has been no exception to this [29], [30], [31]. In their summary article, Karl and Whishaw [9] presented several arguments to show that vision is essential for the integrated control of reach and grasp. Without this feedback information, subjects appear to reach with a grasp aperture which is maximized early to produce results in the face of uncertainty and only refined following contact [32]. In the other direction, slow movement can be considered to be more influenced by feedback [33], [34], [35], in this case, primarily proprioceptive.

We used a Machine Learning or decoding approach in order to probe and characterize the difference in muscular activities of the shoulder and elbow muscles as traditional methods failed to do so. This decoding approach has previously been successfully used to obtain insight into electromyographic activity during gait [36], Whole Body Pointing [37] and Reaching [38]. It provides several advantages when compared with traditional statistics. In the first place, the use of Machine Learning allowed us to study muscle activity in groups. As muscles do not act in isolation, but in groups, this is more in keeping with the physiological manner in which movement is performed. Secondly, muscle electromyographic (EMG) signals are well known for their high variability [39], contributing in good part to motor control studies restricting themselves to the kinematic level. Machine Learning methods with their reliance on stochastic distributions are more adapted to such variability. Our group demonstrated in previous studies that the influence of task variables can sometimes be observed at the level of muscle populations even when they are not significant at the level of single muscles [37], [38], [40], [41]. In this study we also used the approach to provide a dynamic vision of adaptation over the entire time course of reach and grasp. This is in contrast to traditional univariate statistics which provide a more static picture, focusing on gathering information from specific points over the course of a movement.

## Results

In this section we present the results of our study on the patterns of shoulder, elbow and hand muscle modulation as a function of hand orientation. A decoding approach was used to highlight and visualize the differences in muscle adaptation. In our first study, we asked if information concerning hand orientation is present in the shoulder as well as elbow muscle groups. The presence of this information, as reflected in higher than chance predictions of hand orientation, would indicate adaptation of these muscles as a function of hand orientation. In the next section, we used a sliding window over the course of ‘reach’ in order to obtain an idea of the differences in the dynamics of this adaptation. A t-statistic between the EMGs of each muscle in the horizontal versus vertical condition (H-V EMG t-statistic – see Methods) was used to explain some characteristics of the prediction profiles of each muscle group. This section was followed by an investigation of how changes in contexts such as the eyes closed or slow movement conditions affected adaptation in the 3 muscle groups. We finished the Results section by moving from a muscle group analysis, to investigating how much the individual muscles played a role in adapting to hand orientation.

### Elbow and shoulder muscles alter activation as a function of hand orientation

A first step was to look at the entirety of the shoulder, elbow and hand EMGs during reach to see if the decoding algorithm was able to discriminate between horizontal and vertical grasp. The ability to do so would indicate the presence of consistent modifications for target orientation in the individual muscle groups. It would also provide a benchmark against which we could then examine the dynamics of adaptation over the course of the action (see section below).

A first visible result was that all muscle groups had accuracies in identifying hand orientation which were well above chance (above 70%). This means that each muscle group contained some information about the final hand orientation i.e. both the shoulder and elbow muscle groups adapted their contractions as a function of final hand orientation. The adaptation for hand orientation as indicated by decoding accuracy was found to be significantly different for the three muscle groups (one-way ANOVA, *F*(2,27) = 15.985, *p* < 0.001). It was highest for the hand muscle group. This result might appear surprising as the target size did not change and the hand muscles of the group were primarily capable of extension and flexion, rather than rotation. A possible explanation lies in a difference of grasping strategy between horizontal and vertical grasp. Participants tended to extend their wrist during the vertical grasp, resulting in the index on top of the thumb. This led to a small separation between the index and thumb in the antero-posterior axis. In contrast to this, for the horizontal grasp, the wrist was kept in neutral position, with the index in front of the thumb (see Supplementary Figure 10).

### The Dynamics of Adaptation to hand Orientation

The goal of this section was to follow the time course of muscular adaptation and to see if the two components of ‘reach’, the shoulder and elbow muscles, had similar dynamics in the adjustment for hand position. Therefore, instead of using the EMGs from the entire reaching movement, we only used the data from a sliding window for decoding. The window corresponded to 20% of the entire duration for reaching, from the start of reach to the start of grasp. In Figure 5, we can compare the dynamics of adjustments in the various muscle groups. The x-axis for normalized time starts at 0.2 because of the 20% sliding window interval and ends at 1, where grasp starts. Of note in the figure is how most of the adaptation for hand orientation in the first 40% of the movement takes place in the shoulder muscles. The mean decoding in this interval reaches a peak of 68.5%, demonstrating the presence of small amounts of consistent adaptations in the shoulder muscles for hand orientation. The elbow muscles show the lowest levels of adaptation in the interval. Beyond the 40% mark, decoding by the hand muscles starts to increase and comes close to 80% accuracy before the start of grasp. This timing also corresponds to the moment the wrist angle becomes distinguishable between horizontal and vertical movement (see Supplementary Figure 11), indicating that this accuracy increase is linked to wrist rotation. Unlike the shoulder and elbow muscle groups, adaptation by the hand muscle group is simpler, showing a monotonic increase until the start of grasp. In contrast to the shoulder muscles, elbow muscles display peak adjustments for hand orientation much further into the reach movement, between 70 and 80% of the reach movement. Table 1 : Average normalized time to peak accuracy for each muscle and each condition (with standard deviation). shows how the mean peak time for horizontal versus vertical hand orientation adjustment shows a proximo-distal (shoulder-elbow-hand) pattern. The differences in this time point for the shoulder and elbow muscles, also show that despite similarities in kinematics, the shoulder and elbow muscle groups show distinctly different patterns of modulation for hand orientation.

**Table 1:** Average normalized time to peak accuracy for each muscle and each condition (with standard deviation). With the exception of the eyes closed condition, the times to peak followed a proximo-distal pattern.

|  | Shoulder | Elbow | Hand |
| --- | --- | --- | --- |
| Normal | .47(±.26) | .57(±.23) | .78(±.14) |
| Eyes closed | .53(±.25) | .42(±.23) | .64(±.24) |
| Slow | .40(±.17) | .60(±.24) | .76(±.13) |

To obtain a better understanding of the adaptation dynamics of each muscle group, we computed the t-statistic of the EMGs for all the trials and all the subjects in the horizontal versus vertical orientations at each time point from the start of reach to the start of grasp. We will call this variable the H-V EMG t-statistic. This score reflected the differences between the EMGs at each time point for the horizontal versus vertical conditions. The results of this analysis are shown in Figure 6.

Of particular note is the fact that the extensor carpi and extensor digitorum showed the highest H-V EMG t-statistics, hence providing some explanation of the high accuracy of the hand group during classification. These t-statistics also show a steady pattern of increase as the hand moves towards the target, once again providing a possible explanation for the steady increase in orientation prediction by the hand muscle group in Figure 5. As opposed to this, most of the muscles of the shoulder group displayed peaks of H-V EMG t-statistic early in the movement, in keeping with the early peak in accuracy displayed by this group. Finally, the muscles of the elbow group displayed very different behaviors. In particular, the brachioradialis displayed an increase in its H-V EMG t-statistic during the latter half of the movement, concomitant with the rotation of the wrist. This may explain the peak in prediction by the elbow muscle group during the latter half of the movement.

### Patterns of Adaptation are Altered by Eyes Closed and Speed Constraints

The reach-and-grasp movement in daily life is altered by various constraints. Examples of this could be a visual obstruction of the target or speed constraints. We used the same decoding approach which had been used in the previous section to analyse if the three muscle groups changed their adaptation patterns when normal conditions for reach-and-grasp were altered. Figure 7 and Figure 8 show that the adaptations for hand orientation are very different in these conditions. In the eyes closed condition, the most striking thing is the much earlier adaptation of the hand muscles for target orientation. In normal conditions, a 70% accuracy in decoding hand orientation is only achieved after 0.4 of normalized time. In the eyes closed condition however, the modifications for these two hand orientations have already taken place by 0.2 of normalized time. Shoulder and elbow adjustments for hand orientation also appear to be much less consistent, giving rise to lower decoding accuracies than in normal conditions. A comparison of the peak adaptation points of the 3 muscle groups in the 3 conditions can be seen in Table 1 : Average normalized time to peak accuracy for each muscle and each condition (with standard deviation).. The table shows significantly different times to peak for adaptation (as quantified by accuracy) as a function of hand orientation in all 3 muscle groups (two-way ANOVA, *F*(2,81) = 11.784, *p* < 0.001).

In comparison to thef eyes closed condition, the constraint of slow movements produced lower prediction accuracies for the hand muscle group in particular. Another interesting alteration with the change in task constraints was the correlation between the adaptations of the elbow and shoulder muscle groups. It increased in the conditions with imposed constraints. The correlation for average accuracy between the shoulder and elbow muscle groups was -0.36 in normal conditions. This increased to values of 0.64 in the eyes closed condition and 0.79 for slow movements.

### Individual muscles

The next step was to analyze the adaptations of the individual muscles, by examining their capacity to predict hand orientation when isolated in the input vector. As explained in the previous sections, a higher prediction accuracy would indicate consistent muscular adaptations for hand orientation (Figure 8).

The first thing to note was that the accuracy of any individual muscle to predict hand orientation was at least 5% lower than the accuracy of the group to which it belonged (see Figure 4) for a given condition. This demonstrates that the analyses with muscle groups increase the data separability for the two hand conditions. In comparing the adaptations of the individual muscles, the accuracies of the wrist extensors were much higher than those of the wrist flexors, indicating that most of the consistent adaptations for orientation were made during wrist extension for all three conditions.

It is also interesting to observe the shifts in the accuracies of the shoulder and elbow groups for the normal speed versus slow conditions. During movement executed at normal speed, the brachioradialis is the muscle in the elbow and shoulder groups that displays the most differences as a function of hand orientation. However, when movements are slow, its capacity for prediction falls behind those of the other shoulder and elbow muscles, indicating that in slow conditions, the latter muscles play a more consistent role in adaptation, hence indicating an important change in muscle recruitment and strategy for slow movement.

## Discussion

This study aimed to deepen our understanding of the reach and grasp task. While there has been much research on this movement undertaken at the kinematic level [6], [16], [18], [44], [45], [46], [47], [48], there has been much less done at the muscular level. We found that the ‘reach’ phase of a reach-and-grasp movement uncovered very different profiles of its components at the muscular level (shoulder, elbow and hand groups) despite highly correlated activity at the kinematic level. These differences were seen for both the amplitude and temporal aspects of the adaptation profiles.

A first point to be made is that we were only able to uncover the differences in muscular activity as a function of hand orientation using Machine Learning. Univariate statistics did not reveal statistical differences in many of the variables that are usually used in studying motor control such as maximum amplitude or time to peak (Figure 3). This difficulty might have been due to the high variance that is generally found in muscular activity [39], [49]. It might also explain the lower number of studies probing the muscular level for reach and grasp. As a method capable of integrating all alterations in EMG activity, be it change in the time to peak, maximum amplitude, start time or rate of rise, Machine Learning offers a higher capacity for data separation. The method also allows for the analysis of *combinations* of muscle groups, allowing for data separation when any individual muscle may only show weak differences [38].

Figure 5 displays the dynamic profiles for adaptation to orientation of the shoulder, elbow and hand groups. The most visible result from these profiles was that all three muscle groups display very different adaptation profiles for orientation. This is especially interesting for the shoulder and elbow muscles. A large portion of the studies on reach and grasp have been conducted using kinematics and have treated ‘reach’ as one component. This would be justified in many ways by the correlation displayed between these segments [3], [5], [21], although some minor exceptions to this rule have been reported by Desmurget et al [4]. Our results show that while kinematic activity between the joint angles are highly correlated for reach and grasp with different orientations, muscular adaptation for specific conditions is more modular, calling into play some muscle groups more than others. It should be noted however that the lack of correlation for adaptation does not mean that the EMG amplitudes of the different arm segments were not correlated. Amplitudes can change to adapt the arm and hands for different orientations while still being correlated.

Differences in the adaptation profiles of the three muscle groups were seen in their temporal patterns as well as amplitude. Concerning temporal patterns, the hand adaptation profile is relatively simple. In contrast to this, both the shoulder and elbow profiles show multiple peaks. One mechanism leading to this difference could be the contrast between the balance of feedforward and feedback control for these muscle groups. A more complex profile may be the result of multiple controllers such as a combination of feedforward and feedback control, while a simpler profile might indicate more feedforward regulation. A possible mechanism of adaptation in this scenario might be a greater role for feedforward control in the case of the hand muscles, while the correction and fine-tuning for final corrections take place in the elbow muscles and shoulder muscles which both show a second decoding peak close to grasp onset (normalized time point 1). Extensive future studies would be required to investigate this possibility. Another thing to be noted in these profiles is that adaptation of the muscle groups responsible for hand orientation occurred well before contact with the target. This is in agreement with the studies by Desmurget et al. [3], [4] who observed an almost linear relationship between the wrist displacement and its rotation angle during reach. Indeed, several previous studies on grasp have shown that hand pre-shaping for prehension occurs well before contact with the target [14] and that it is adjusted for target features like size [8], [44].

Another clear difference between the groups was in terms of amplitude. Classification accuracy was clearly higher for the hand muscles in comparison to the shoulder and elbow muscles. This indicates that the most consistent adaptation for hand orientation came from the hand muscles. This is surprising given that these muscles are not key in creating hand rotation, but were rather composed of extensor and flexor muscles. The results suggest that the manner in which the same target was gripped was altered by orientation. The contribution of other muscles that do not play a key role in rotation to the adaption for object orientation can also be seen at the shoulder and elbow levels in Figure 9. The higher than chance prediction capacities of muscles like the Triceps or the Extensor Digitorum which do not play important roles in arm or hand rotation, shows that the adjustments for grip orientation did not only involve rotation, but in the transport of the arm to the object. This is similar to the manner in which some researchers have shown that the kinematics of transport are altered by the nature of the task to be performed by the hand, such as point, hit or grasp [50], or by Desmurget et al [3], [5] who described how wrist path and trajectories were highly altered by object orientation.

Despite their different adaptation profiles, all the muscle groups had one thing in common. They all showed adaptation for target orientation very early at the start of movement. This is seen by classification accuracies that are significantly above chance at the onset of reach (normalized time point 0.2). It may therefore be the result of feedforward adaptation for orientation in all three muscle groups. Feedforward muscular activity is generally considered as what takes place within about 80ms after the start of movement [51], [52]. To go further, the initial decoding capacities of all three muscle groups in Figure 5, may even be part of the Anticipative Postural Adaptations (APA) which are seen even before the onset of movement itself. Several researchers have now shown that APAs are modulated as a function of task conditions for a variety of movements [37], [53], [54]. Once again, this question can only be resolved in future studies in which we check if the Machine Learning algorithm is capable of identifying hand orientation using a short EMG segment *before* the onset of movement.

Alterations in sensory feedback had very strong effects on the modulation profiles of the three muscle groups. In both the reduced feedback condition without vision and the increased feedback condition of slow movements, classification accuracies were reduced for the shoulder and elbow muscles. This need not necessarily mean that they no longer contribute to altering hand orientation. As in the case of any statistical method, it could be due to a higher variability in their contribution. The shifts in the times to peak of the contribution of hand muscles is also interesting. The reduced vision condition leads to the earlier onset of hand muscle determination. This result can be compared to the study by Karl et al [32] where reduced vision led to subjects opening their hands early but refining the details of grasp upon contact with the target. This shows that the onset of hand muscle contribution in the case of reduced feedback is variable and context dependent. The results with the slow movements in our study on the other hand produced a delayed onset in hand muscle contribution to hand orientation. In both cases with altered feedback, there was an important increase in the correlation of adaptation by the shoulder and elbow muscle group (Figure 7 and Figure 8). This apparent reduction in the number of degrees of freedom in the system is reminiscent of the Gimbal lock situation in rotating systems in which there is a loss of some degrees of freedom [55], [56]. A final point to be made concerning the situations with altered feedback is that despite the changes in muscle adaptation profiles, they all showed a similar level of decoding capacity at the start of the movement, as in normal conditions. This supports the idea that the early decoding levels are indeed the result of feedforward activities and hence are unaltered by changes in feedback.

In conclusion, despite their correlated activities at the kinematic level, the shoulder, elbow and hand muscles show very different dynamics in adapting to hand orientation. This is particularly interesting in the case of the shoulder and elbow segments which are usually considered as a unified whole for ‘reach’. All the muscle groups showed a very early adaptation as a function of orientation. The time to peak for adaptation of the three muscle groups displayed the ubiquitous proximal-distal pattern, starting with the shoulder, followed by elbow and then hand muscle groups. Alterations in sensory feedback were found to strongly influence the dynamics of muscle activation patterns. We suggest that Machine Learning, by undertaking a multivariate analysis, can be a complementary method for studying the dynamics of motor control.

## Methods

### Experimental setup and task

Eighteen participants (6 men, 12 women, average age of 20.6±2 years) were invited to the laboratory. The general procedure of the experiment was explained to them and they were allowed to ask the experimenter questions except those concerning the hypothesis of the experiment. This was done in order to keep participant movements as natural as possible. Then, they signed a consent form for the experiment and filled out a demographic questionnaire and the Edinburgh Handedness Questionnaire. The experiment was conducted following the guidelines of the Helsinki declaration and the ethics committee of INSERM (Institut National de la Santé et de la Recherche Médicale) number C03-07-29-06-2007 approved the experimental protocol.

The participants were then equipped with wireless EMG sensors (PicoEMG, Cometa, Italy) on twelve muscles of the shoulder, arm and forearm (see Figure 1). The recorded muscles were : the pectoralis major (PM), the posterior deltoid (PD), the medial deltoid (MD), the anterior deltoid (AD), the biceps (Bi), the short triceps (ST), the long triceps (LT), the brachioradialis (Br), the extensor carpi (EC), the flexor carpi (FC), the extensor digitorum (ED) and the flexor digitorum (FD). Reflective markers were also placed on the shoulder, elbow, wrist, thumb, and index finger of the participants. Their positions were followed by a position tracking system (Vicon Motion Systems, UK). Finally, to investigate the loss of visual feedback, participants were equipped with visual occlusion spectacles (PLATO, Translucent Technologies Inc., Canada).

**Figure 1:**
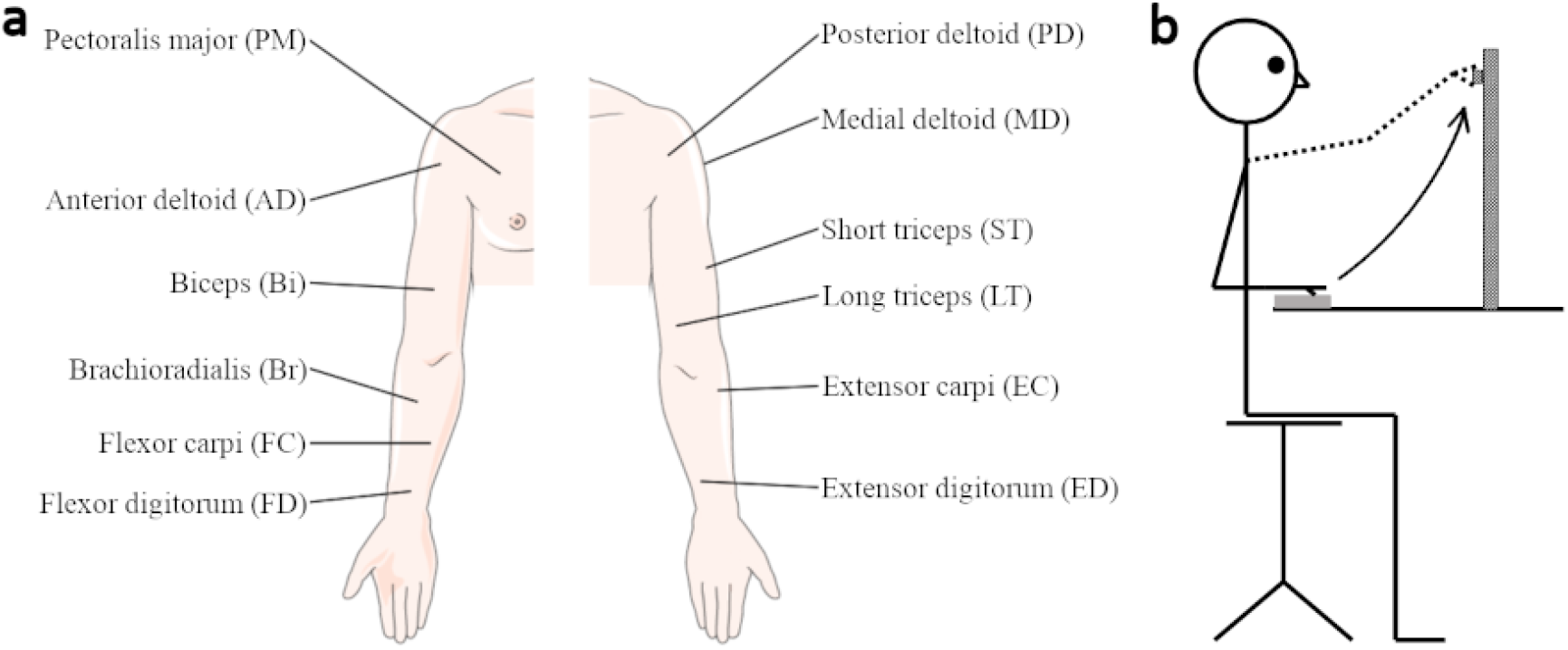
a) Schematic of the EMG electrodes placement and the muscles targeted. B) Illustration of the reach-and-grasp task : Participants started with there hand resting on a plate then move to grasp the target placed in front of them.

Participants sat in front of the experimental apparatus consisting of a tall vertical bar (45cm) to which was attached a small bar (3×10cm). The small bar could be swiveled so that it had a horizontal or vertical orientation. Reflective markers attached to the apparatus allowed us to follow the distance of the hand from the target. A cushion that served as a starting point for the participants was also placed on the same table on which the experimental apparatus was placed.

Participants were asked to keep their hand relaxed but extended and flat on the cushion at the start of each trial and to come back to this position at the end of the trial. They had to reach for the target, grasp it between their thumb and index finger, and then bring it to the cushion where they could rest their hand. They were asked to particularly pay attention to grasping the target with a vertical or horizontal pincer grasp depending on the orientation of the rotating part (see Figure 1.b for a diagram of the apparatus and the experiment).

Participants performed the movement at either a natural speed (“about 3s for the complete movement”) or at an exaggeratedly slow speed (“about 15s for the complete movement”). They were verbally notified by the experimenter of the expected speed before each trial and received feedback on their execution of the task. Moreover, the occlusion spectacles were opened or closed under the control of the experimenter, with the condition for the next trial being set at least 1s before the GO signal for the trial was given.

Participants did 15 trials per condition for a total of 120 movements (2 *orientations* × 2 *speeds* × 2 *eye conditions* × 15 *trials*). Trials were divided into 2 blocks, one for horizontal movements and the other for vertical movements. The first block to be performed was randomized. The other conditions were randomized within the block. Participants took breaks every 30 movements to avoid tiredness.

After the experiment, participants were debriefed and the study aim and hypothesis were explained to them.

### Preprocessing

Each trial was cut using the wrist marker movement: the start of the movement was defined as when the wrist marker moved 5mm away from its resting position and the end of the movement was defined when the wrist position reached a maximum in the sagittal plane. The EMG signals between these two events were filtered (5^th^ order Butterworth filter, bandpass between 30 and 300 Hz) then rectified and integrated over a sliding window of 100ms. After that, the EMGs were all normalized in time. Finally, the recordings were normalized by the maximum amplitude for each muscle and each participant separately. An example of recording for one participant can be seen in Figure 2.

**Figure 2:**
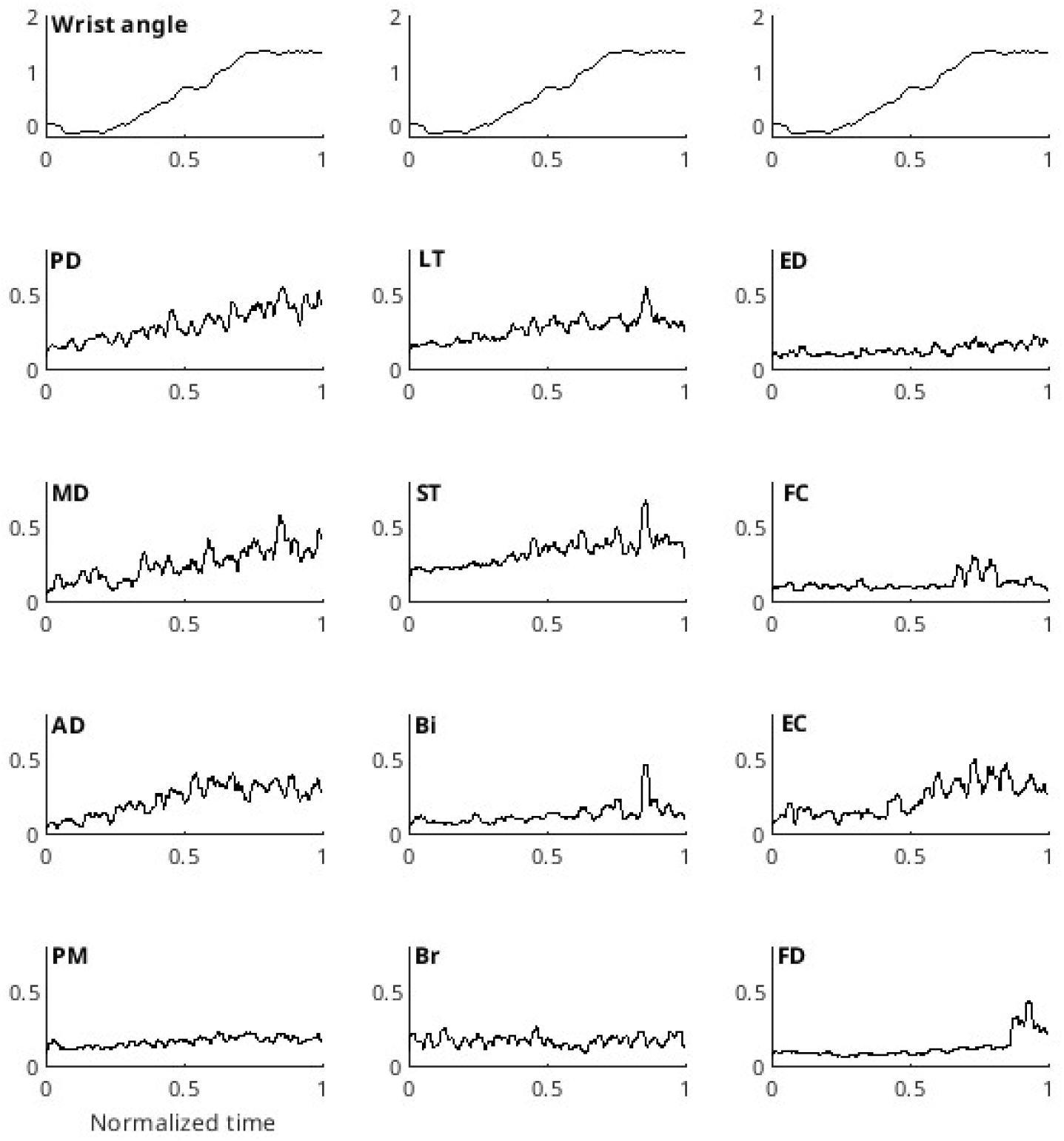
RMS of the muscular activity during one vertical grasp trial of a subject for each recorded muscle. The first line of the graph displays the wrist angle (see Methods). It is to be noted that the three graphs along this line are identical and have been repeated to facilitate interpretation of the RMS data in each column.

The wrist angle was computed as the frontal angle (elevation angle) between the thumb-index vector and the horizontal at each time step and then averaged for all trials of a participant in a given condition.

### Machine learning processing

We used a Machine Learning approach in this study which allowed us to draw simultaneously from several differences in the EMG such as time shifts and amplitude changes. Figure 3 shows that the use of individual variables, as is typically done in univariate statistics, did not yield any significant differences between the muscles analyzed for the two hand orientations. We had compared the maximum amplitude and the time to peak for each muscle in horizontal or vertical conditions. These two values were chosen as they are classical measures of muscular activity. Indeed, when using classical univariate analysis, a specific measure has to be extracted from the data, potentially losing information in the process and painting an incomplete picture of the changes. A Two-way ANOVA (Muscle x Grasp orientation) was conducted for each measure. In both cases, the grasp orientation effect was not significant : F(1,408)=2.991, p=0.08 for the maximum amplitude and F(1,408)=0.975, p=0.32 for the time to peak. Therefore, if this univariate analysis had been conducted alone, we would have concluded that there was no significant difference in muscular activity between horizontal and vertical movements. However, the machine learning algorithm was able to discriminate the muscular activity between those two conditions with an accuracy of up to 85% (see Figure 4), highlighting the failure of univariate statistics for performing the task.

**Figure 3:**
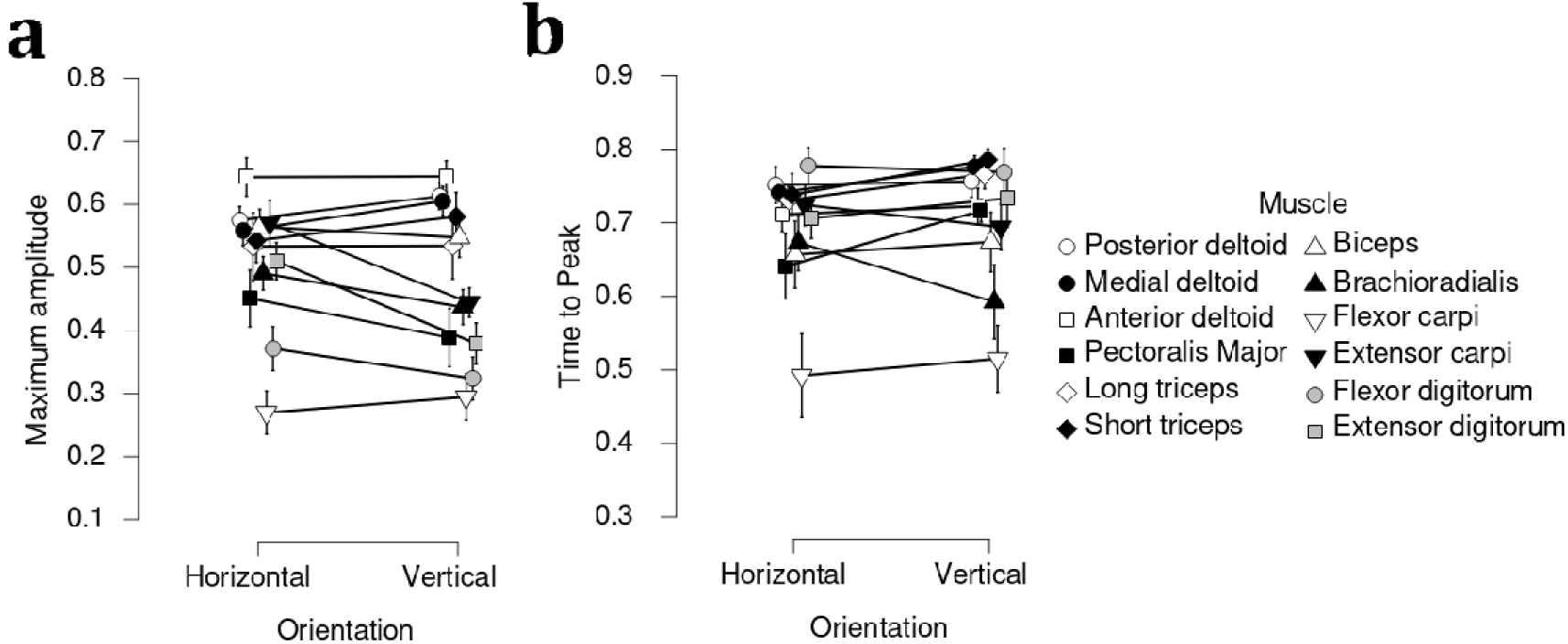
a) Maximum amplitude and b) Time to peak for each muscle in normal conditions. Error bars indicate standard error. No significant differences were observed in any muscle for these variables between Horizontal and Vertical orientations.

**Figure 4:**
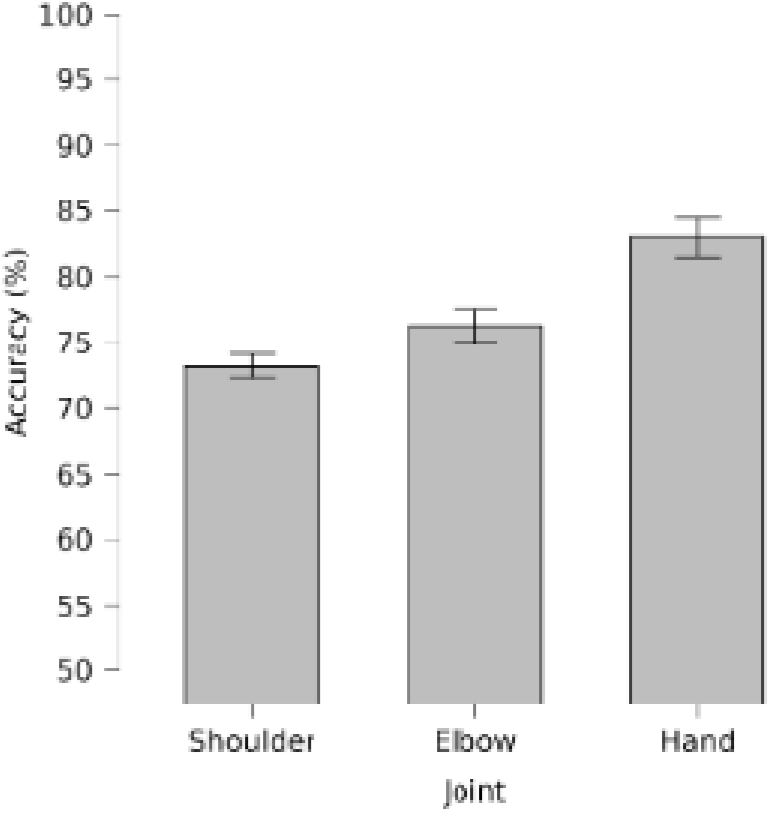
Accuracy of algorithm discriminating between horizontal and vertical movement during normal conditions for each muscle group. Error bars indicate standard deviation. The figure shows that all three muscle groups had different activities as a function of hand orientation. The accuracies between the three groups were found to be significantly different.

**Figure 5:**
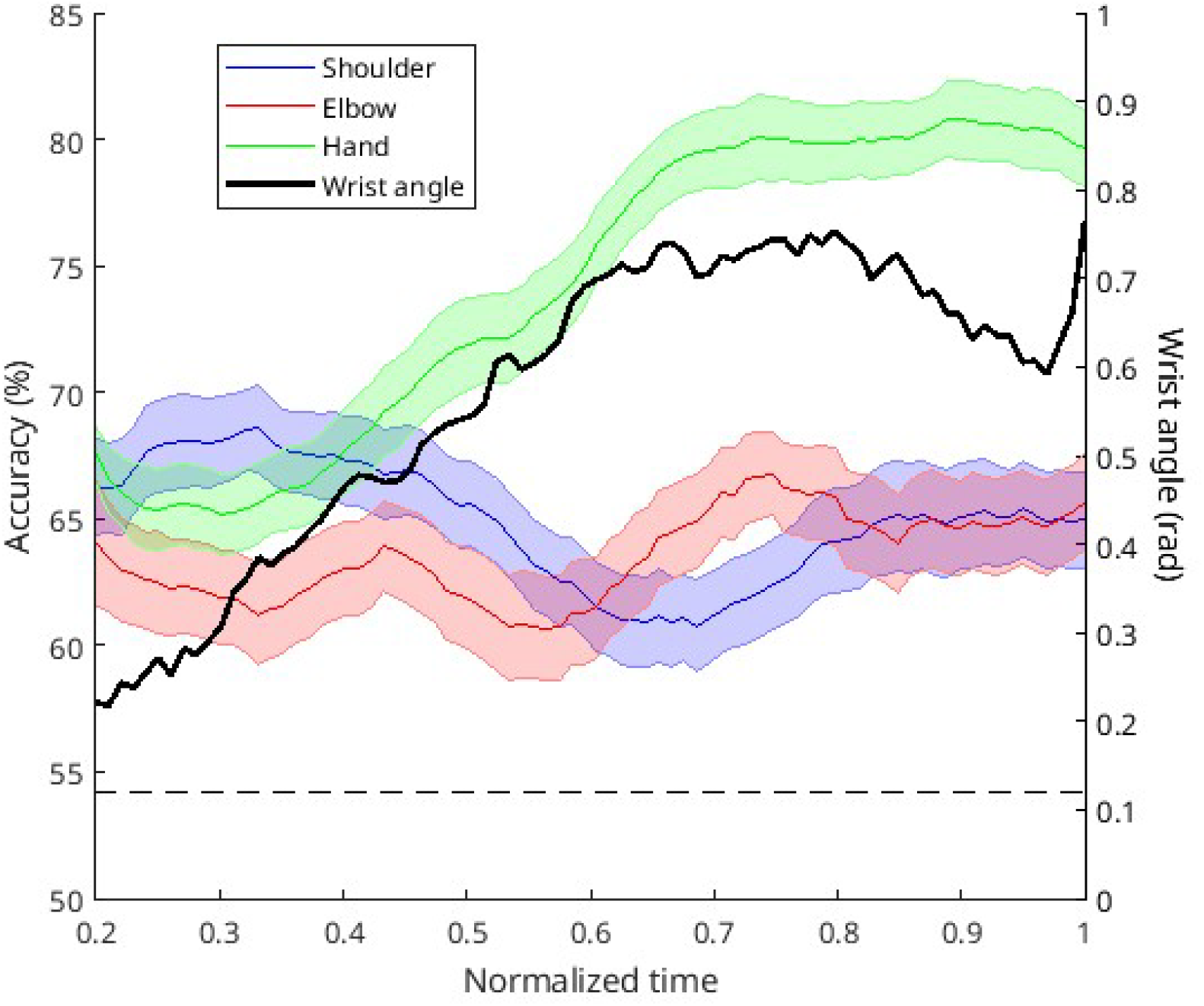
Time evolution of the discriminating accuracy between horizontal and vertical reach for each muscle group in normal condition. Shaded areas indicate standard deviation. The dashed black line indicates the minimum accuracy to exclude random classification. The black full line of the graph displays the wrist rotation angle.

**Figure 6:**
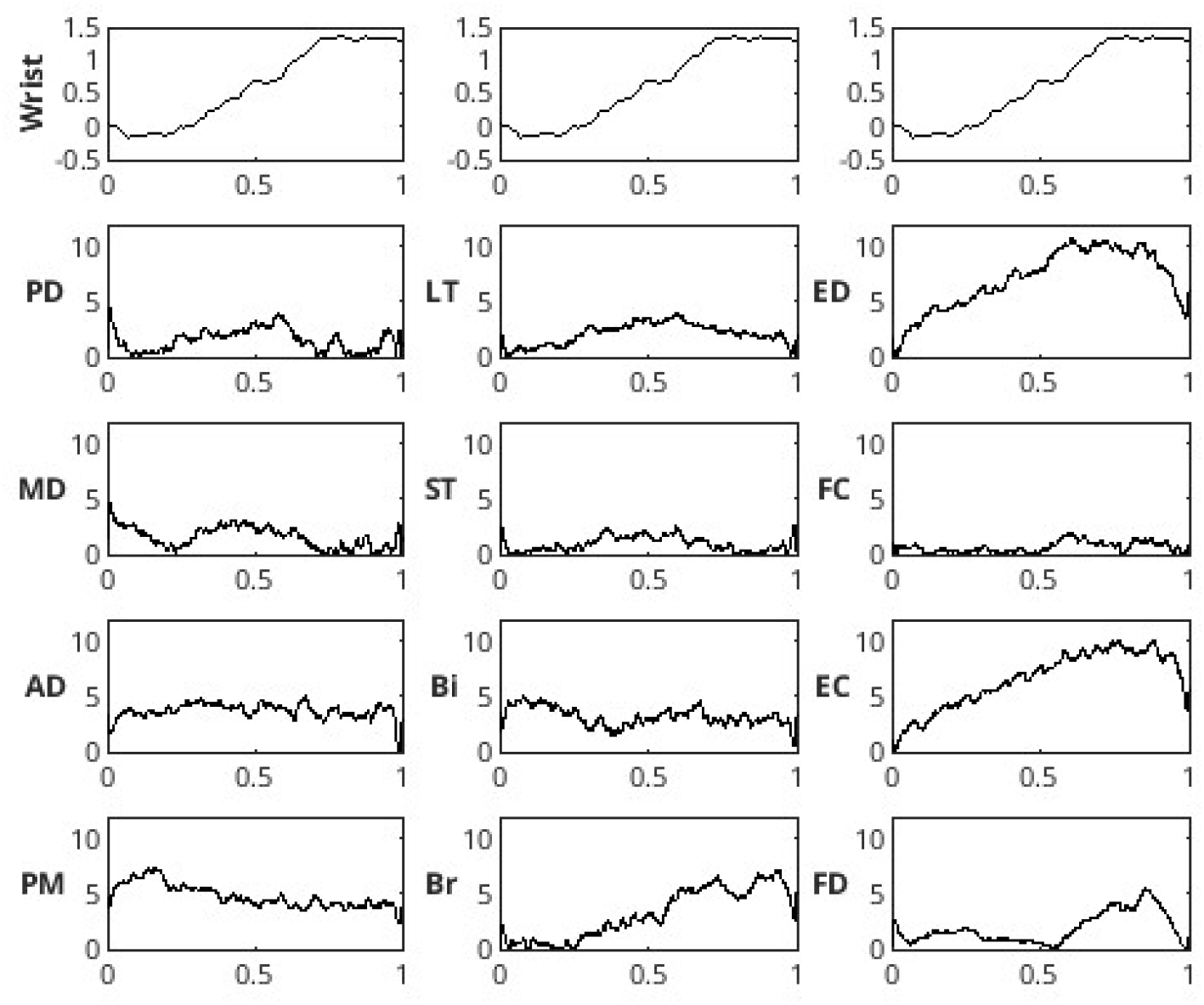
The H-V EMG t-statistic for each time point of the EMGs for all trials and all subjects in the normal condition. The score reflects the separation between the EMGs of the horizontal versus vertical movement at a given time point. The first line of the graph displays the wrist rotation angle for one trial taken as an example. It is to be noted that the three graphs along this line are identical and have been repeated to facilitate interpretation of the H-V EMG t-statistic in each column.

**Figure 7:**
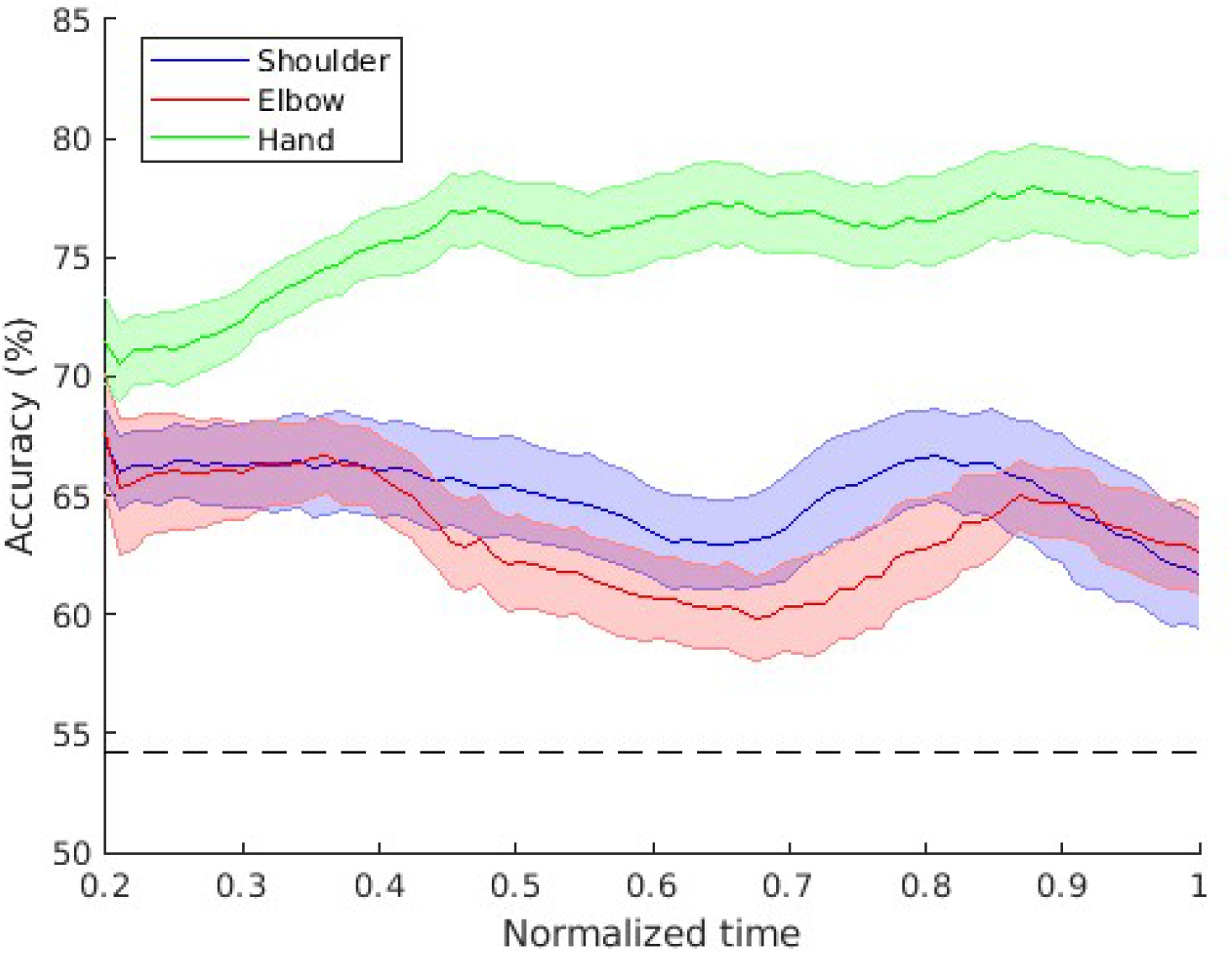
Time evolution of the discriminating accuracy between horizontal and vertical reach for each muscle group in the eyes closed condition. Shaded areas indicate standard deviation. The dashed black line indicates the minimum accuracy to exclude random classification.

**Figure 8:**
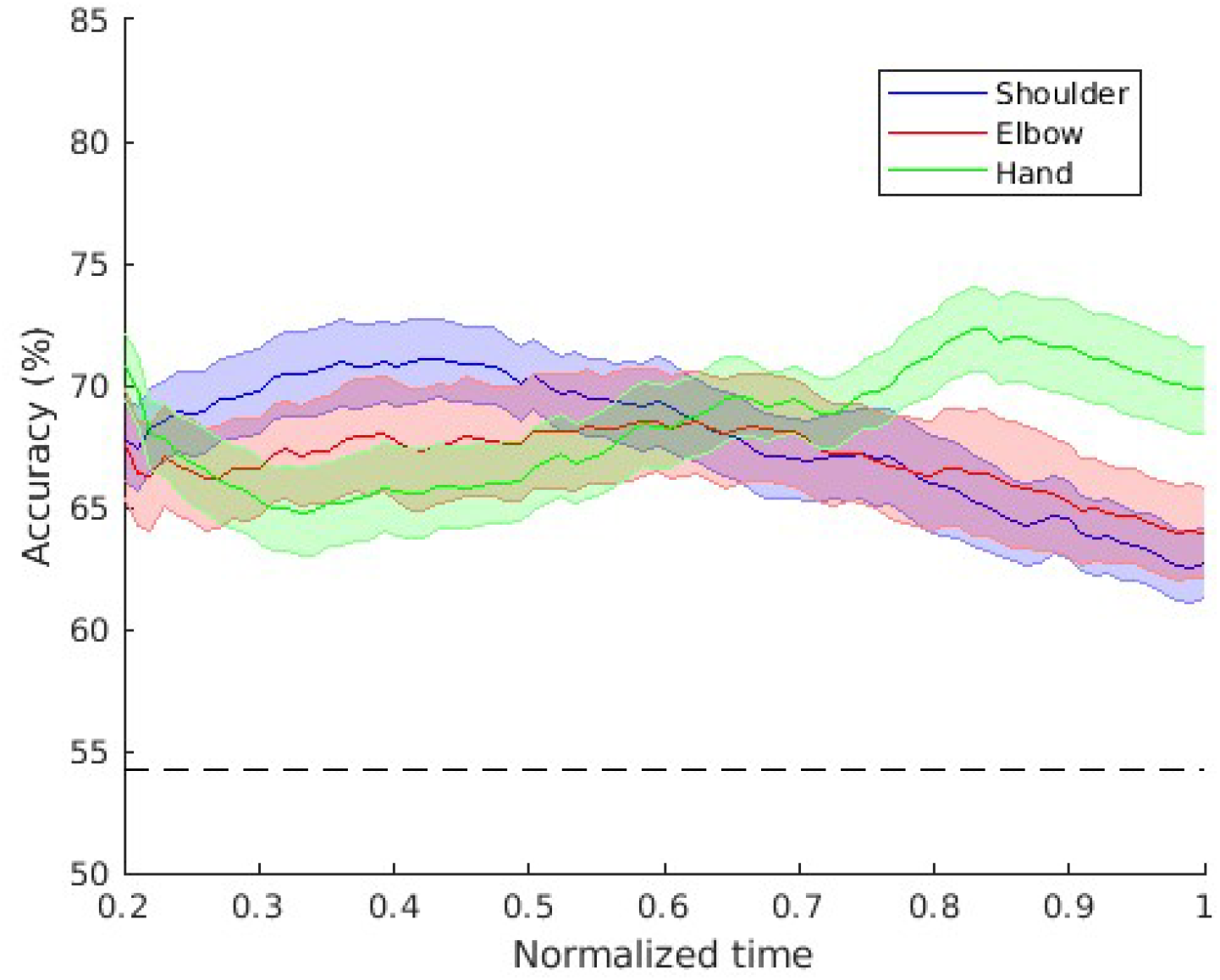
Time evolution of the discriminating accuracy between horizontal and vertical reach for each muscle group in the slow movement condition. Shaded areas indicate standard deviation. The dashed black line indicates the minimum accuracy to exclude random classification.

**Figure 9:**
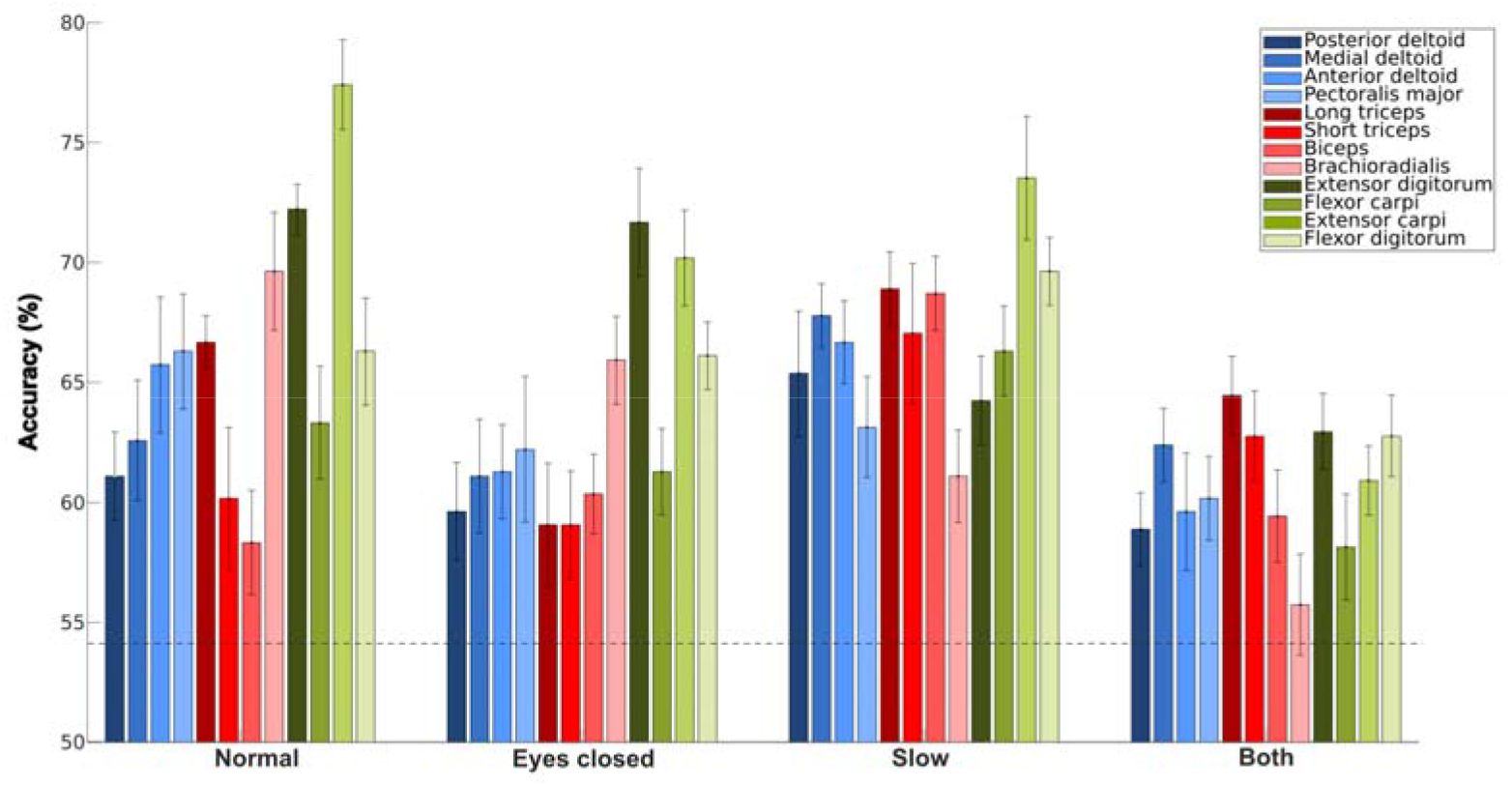
Accuracy of algorithm discriminating between horizontal and vertical movement for each muscle and each condition. Error bars indicate standard deviation. The dashed line indicates the minimal accuracy required to reach a significant level of discrimination.

The use of Machine Learning also allowed us to put muscles into anatomical groups, helping us to gain comprehension with a big dataset. We used a binary classification approach to discriminate between Horizontal and Vertical pinch during the grasp based on part or complete EMG traces. The logic behind this paradigm is that if the algorithm is able to discriminate between the two pinch orientations solely based on the EMG recordings, those must contain some specific differences in the muscular activation pattern between the two orientations. To account for the risk of overfitting, we used a 10-fold cross-validation setup in all analysis.

Moreover, to obtain a better understanding of the important features for the algorithm, we computed the t-statistic between the Horizontal and Vertical grasp EMGs at each normalized time point and for each muscle. This measure is useful as it gives a partial indication of the separability of the data but fails to account for interaction between features.

The algorithm used was ADABoost. The Adaptative Boosting algorithm, commonly known as ADABoost, was created in 1997 by Freund and Shapire [42]. This algorithm functions under the principle of boosting : a series of weak learners are trained in succession, each new learner aiming to have a good accuracy on the data the previous one failed to correctly classify. When presenting new data, the decisions of all learners are weighted to obtain the final decision of the algorithm. As long as the weak learners have above chance accuracy, the complete algorithm will be a strong classifier.

To test different hypotheses, either complete or partial EMG traces were fed to the algorithm for training (and testing). More specifically, muscles were separated into 3 groups based on the main joints they have an action on [43] :

- Shoulder : Pectoralis major, Anterior deltoid, Medial deltoid, Posterior deltoid
- Elbow : Long triceps, Short triceps, Biceps, Brachioradialis
- Hand : Extensor carpi, Flexor carpi, Extensor digitorum, Flexor digitorum

Thus, depending on the analysis, the algorithm potentially only had access to the EMG data from one of these groups. Finally, one type of analysis, the sliding window analysis, only provided the data to the algorithm from a sliding window of 20% of the duration of the movement.

Accuracy in correctly identifying whether the provided test EMG segment came from a horizontal or vertical grasp gave us a quantitative measure of data separation. A high accuracy above chance levels (50%) would indicate good EMG separation between the horizontal and vertical conditions. To account for chance, we computed the cutoff at which the accuracy would deviate from pure chance with a level of 95% confidence. For the number of trials in our experiment (18 × 30 for each condition since we are using a cross-validation), the cutoff to exclude the possibility that the classifier was purely random was 54.2% accuracy.

## Acknowledgement

The authors want to thank Pr.Jérémie Gaveau for his valuable inputs while starting this study.

## Authors contributions

FC participated in the conception of the study, collected the data, analyzed it and participated in preparing the paper.

PH and NC participated in the interpretation of the data analysis and paper preparation. ET participated in the conception of the study, the interpretation of the data analysis and paper preparation

## Data availability statement

The datasets generated and analysed during the current study are available from the corresponding author on reasonable request.

## Supplementary figures

**Supplementary Figure 10:**
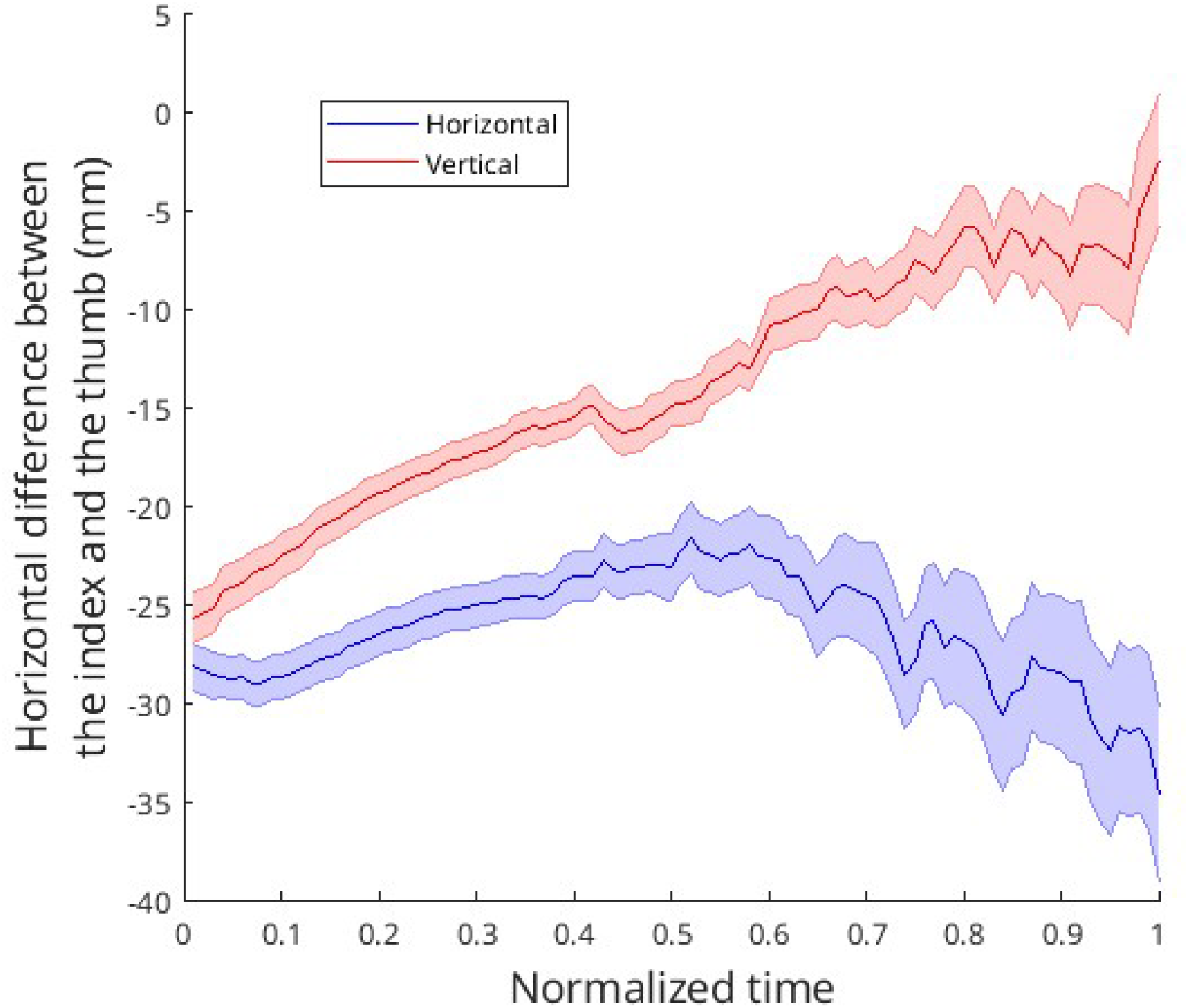
Distance between the index and the thumb along the antero-posterior axis of the hand for horizontal and vertical grasp. Shaded areas indicate standard deviation among all participants.

**Supplementary Figure 11:**
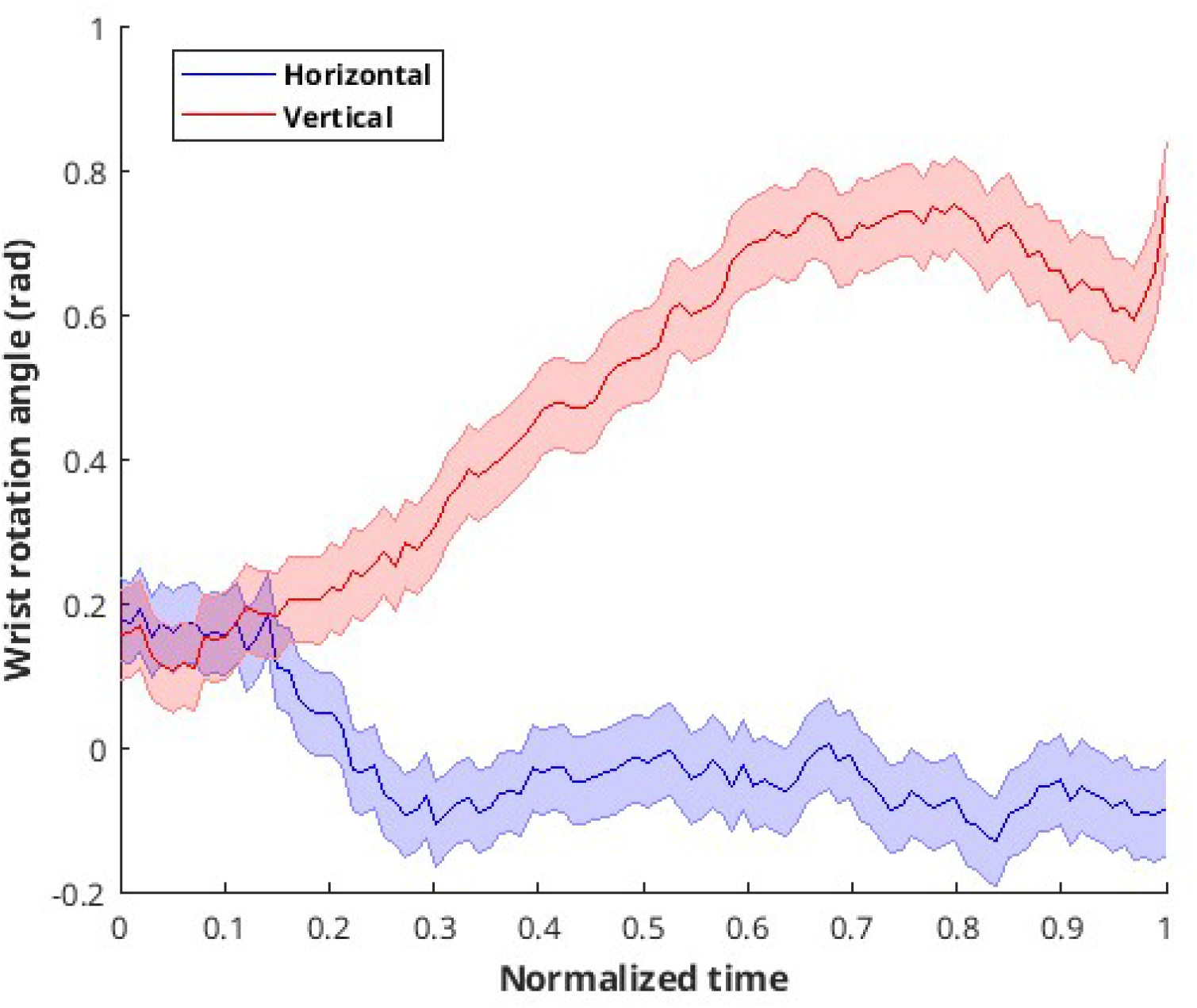
Frontal angle between the index-thumb vector and the horizontal for horizontal and vertical grasp. Shaded areas indicate standard deviation for one participant.

